# Molecular evidence for the hybrid origin of *Cryptocoryne ×purpurea* Ridl. nothovar. *purpurea* (Araceae)

**DOI:** 10.1101/2020.09.09.289165

**Authors:** Rusly Rosazlina, Niels Jacobsen, Marian Ørgaard, Ahmad Sofiman Othman

## Abstract

Natural hybridization has been considered a source of taxonomic complexity in *Cryptocoryne*. A combined study of DNA sequencing data from internal transcribed spacer (ITS) of nuclear ribosomal DNA and *trnK-matK* region of chloroplast DNA was used to identify the parents of *Cryptocoryne* putative hybrids from Peninsular Malaysia. Based on the morphological intermediary and sympatric distribution, the plants were tentatively identified as the hybrid *Cryptocoryne ×purpurea* nothovar. *purpurea;* plants were pollen sterile and had long been considered to be hybrids, possibly between two related and co-existing species, *C. cordata* var*. cordata* and *C. griffithii*. The *C. ×purpurea* nothovar. *purpurea* status was independently confirmed by the presence of an additive ITS sequence pattern from these two parental species in hybrid individuals. Analysis of the chloroplast *trnK-matK* sequences showed that the hybridization is bidirectional with the putative hybrids sharing identical sequences from *C. cordata* var*. cordata* and *C. griffithii*, indicating that both putative parental species had been the maternal parent in different accessions.

## Introduction

Natural interspecific hybridization has been demonstrated to be an important force in forming new species [1, 2] and plays a crucial role in plant evolution and diversification [3, 4]. The occurrence of natural hybridization between different species, however, is not universal but concentrated in a limited fraction of plant families and genera [5]. Natural hybridization has been suggested to occur frequently in *Cryptocoryne*. The genus can be seen as having multiple populations in various river systems and hybridization can be an evolutionary driving force that constantly creates new genotypes that are spread across the ever-changing river systems [6–9]. Most recently, 64 species, 19 varieties and 14 interspecific hybrids have been recognized [9–13]. Recognizing the *Cryptocoryne* hybrids began in the 1970s [14–16] and two of the *Cryptocoryne* hybrids have been recognized (as species) for more than 100 years, it was not until after 1975 that it was realized that some of the plants were probably interspecific hybrids [14]. The uncertain status and tendency of *Cryptocoryne* to hybridize naturally may create more complexity in terms of taxonomic studies and classification. The natural *Cryptocoryne* hybrids have been previously reported from Peninsular Malaysia [6, 14], Sri Lanka [16–18], Thailand and Lao P. D. R. [7, 15], Singapore [19], Sarawak [20], Kalimantan [21, 22] and Sumatera [12], with an overview presented by Jacobsen et al. [7]. Even though *Cryptocoryne* hybrids have much reduced fertility, the hybrids are highly successful due to the proliferous propagation by numerous, long, subterranean stolons, resulting in very large stands of hybrid plants, thus easily detectable in nature.

*Cryptocoryne* × *purpurea* Ridl. nothovar. *purpurea* is a natural hybrid, which can be found in Peninsular Malaysia. According to Othman et al. [6], this plant was first collected from Kota Tinggi, Johor by Ridley in 1892, then was cultivated in the Botanical Garden in Singapore and shipped to Europe in 1898. It flowered at Kew and was pictured as *C. griffithii* Schott in Botanical Magazine in 1900 (t 7719). In 1904, Ridley described this plant as a new species, named as *C. purpurea*. It was cultivated widely as an aquarium plant in Europe in the following years, although it almost disappeared towards the end of the century. Based on the low pollen fertility [14], it was suggested that *C. purpurea* was a hybrid between *C. cordata* Griff. var. *cordata* and *C. griffithii* Schott based on the coherence of morphological characteristics (broad collar zone – *C. cordata*, and purple, rough spathe limb - *C. griffithii*) [6]. de Wit [23] gave a comprehensive explanation of the differences between *C. griffithii, C. cordata* var. *cordata* and *C*. × *purpurea* nothovar. *purpurea*. Evidence for this morphological assumption gained support over the years and it is now generally accepted as a hybrid between the diploid *C. cordata* var. *cordata* and *C. griffithii* and moreover both parents and the putative hybrid are found in the same region [6, 7]. In a study on artificial hybridization between species of *Cryptocoryne* from Peninsular Malaysia, support was presented for *C. ×purpurea* nothovar. *purpurea* being a natural hybrid between *C. cordata* var. *cordata* and *C. griffithii* [8]. Verifying the hybrid origins of taxa in question is valuable for studies on taxonomy, evolution and conservation. If parental species overlap at multiple locations in geographical distribution, hybrids may emerge independently from hybridization between local parental genotype populations and may show obvious morphological differences [24]. Therefore, different hybrid taxa with slight morphological dissimilarity may have evolved as a result of hybridization of the same parental species at separate locations. In the present study, we compared two diverse *C. ×purpurea* nothovar. *purpurea* populations, in streams only 2-3 km apart, which differ in the shape, surface structure and colour of the spathe limb representing two hybridization events with different parental genotypes in the state of Malacca [7]. Therefore, inferring the origins of such hybrid taxa based on morphology alone may be difficult because morphologically similar hybrids can arise from hybridization between different populations of the same parental species or be influenced by environmental conditions; thus, can be unreliable and misleading. In such cases, molecular means have been proven successful in identifying hybrid genotypes and determining the origins of various hybrid taxa [24–27].

To our knowledge, no published reports are available on molecular evidence for the hybrid origin of *Cryptocoryne* species and the role of natural hybridization in the evolution of this genus remains elusive. Combined nuclear and plastid DNA markers provide potentially complementary evidence about a putative hybrid, allowing different questions to be investigated. To verify the hybrid origin of these intermediate individuals in the present study, nuclear ribosomal DNA (nrDNA) internal transcribed spacer (ITS) region was first applied to examine interspecific divergence between the putative parental species and to assess the hybrid origin of the intermediate individuals by the additive patterns of both parental species. Secondly, chloroplast *trn*K-*mat*K region was applied due to their success in evaluating interspecific variation in most angiosperms and also their use in identifying the maternal origin of hybrids.

## Materials and methods

### Plant material

The individuals of the putative hybrid *C. × purpurea* nothovar. *purpurea* and the presumed parental species *C. cordata* var*. cordata* and *C. griffithii* were collected from different locations in order to detect potential intraspecific sequence of polymorphism. We sampled seven individuals of *C*. × *purpurea* nothovar. *purpurea*, five individuals of *C. cordata* var*. cordata* and five individuals of *C. griffithii* respectively. Since interspecific hybridization may be confounded by incomplete lineage sorting among closely related species, additional *Cryptocoryne* species (*C. nurii* Furt. var. *nurii* and *C. schulzei* De Wit) was further examined. All accessions are summarized in Table 1. The geographical distributions of *C. ×purpurea* nothovar. *purpurea* and the putative parental species was provided in S1 Fig.

**Table 1.**
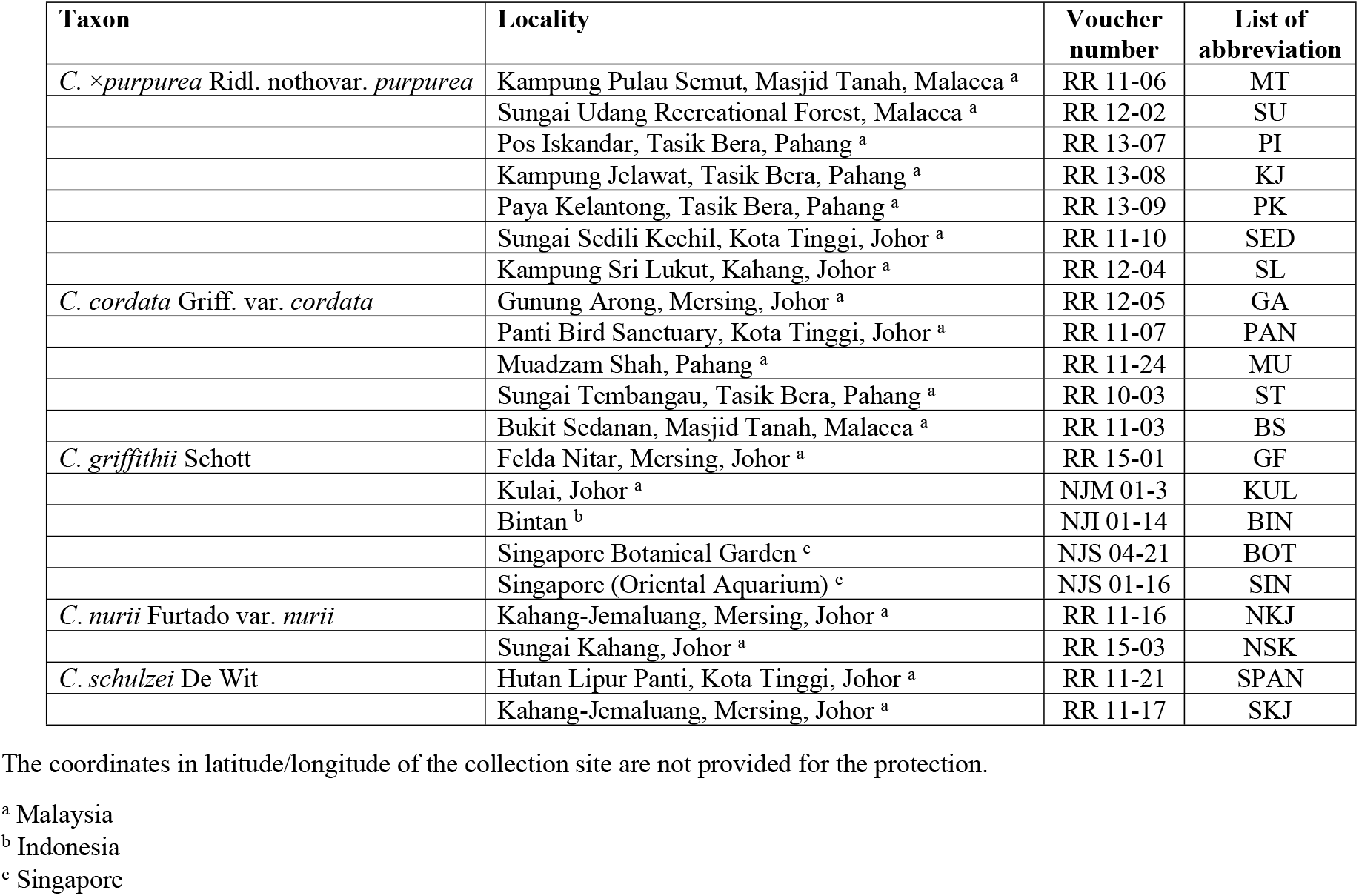
Taxa, localities, voucher numbers and list of abbreviation for sampled *Cryptocoryne* specimens.

Fig 1 showed the spathe limbs of different accessions of *C. ×purpurea* nothovar. *purpurea* and the putative parental species. Voucher specimens have been deposited in the Herbarium Unit, Universiti Sains Malaysia (Penang, Malaysia) and the Botanical Museum, Copenhagen (C) (Natural History Museum of Denmark). Young leaves were cleaned with sterile distilled water before drying with silica gel. Upon completion of the drying process, samples were stored at - 20 °C before being used for DNA extraction.

**Fig 1.**
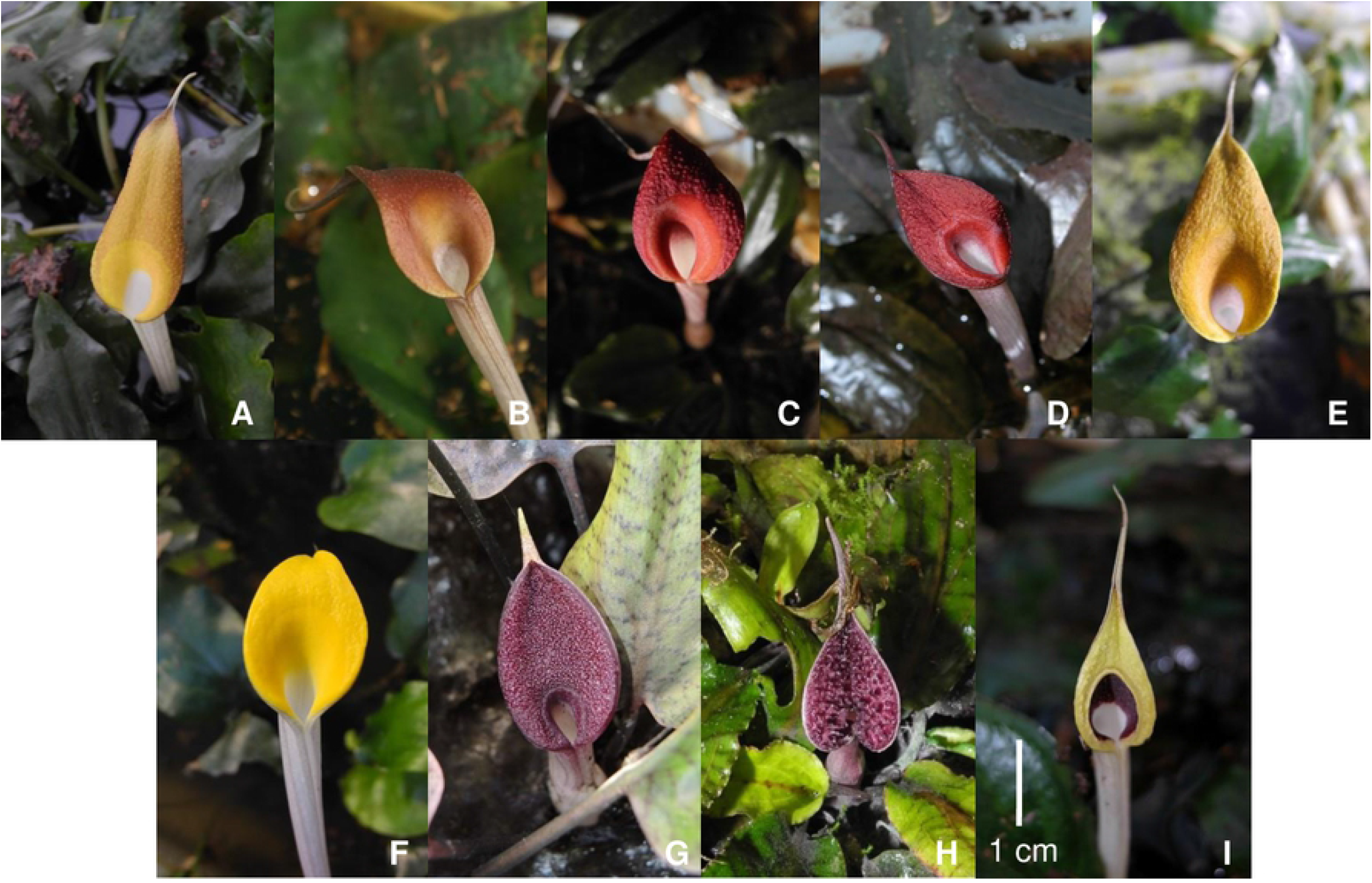
The spathe limbs of different accessions of *C. ×purpurea* nothovar. *purpurea* and the putative parental species. *C. ×purpurea* nothovar. *purpurea* (A, MT; B, SU; C, PI; D, SL; E, SED). *C. cordata* var*. cordata* (F, BS). *C. griffithii* (G, GF). *C*. *nurii* var. *nurii* (H, NSK). *C*. *schulzei* (I, SPAN).

### DNA extraction, PCR amplification and sequencing

Total genomic DNA was extracted from silica-dried leaf tissue using the CTAB protocol described by Doyle and Doyle [28]. The nrDNA ITS and cpDNA *trn*K-*mat*K regions were amplified using universal ITS1 and ITS4 primers [29] and *trn*K-3914F, *trn*K-2R primers [30] respectively. We obtained two newly designed primers for *trn*K*-mat*K regions specific for *Cryptocoryne* species; *mat*KC-450F (5’-AGGGCAGAGTAGAGATGGATG-3’) and *mat*KC-537R (5’-TATCAGAATCCGGCAAATCG-3’). The PCR products were directly sequenced using PCR primers after purification with Gel Clean-up System (Promega Corporation, Madison, WI, USA). All sequencing reactions were done on ABI 3700 DNA automated sequencer with the BigDye chemistry (Applied Biosystems, Foster City, CA, USA). The ITS PCR products which produced unreadable sequence data with superimposed peaks in the chromatograms were further purified using the Wizard SV DNA Clean-Up System (Promega Corp., Madison, WI, USA), and then cloned to ensure representative amplification of the parental copies using pGEM-T Easy-cloning Vector Kit (Promega Corp., Madison, WI, USA) and transformed into competent *Escherichia coli* JM109 at 42 °C. The transformed bacteria were screened on solid LB media with 100 mg/mL ampicillin at 37 °C overnight. The six positive clones with the correct size inserts were confirmed using colony PCR and subsequently sequenced using the ITS primers earlier described. Also, the possibility of artificial recombinants being produced under standard PCR conditions was tested using ITS primers and a 1:1 mixture of *C. cordata* var*. cordata* (GA) and *C. griffithii* (KUL) DNA as template. All of the sequences were deposited in GenBank with accession numbers KU196170-KU196197 and KU196237-KU196248.

### Sequence alignment and phylogenetic analyses

DNA sequences were assembled and aligned using (MEGA) 7.0 [31]. A most parsimonious (MP) unrooted tree was first built for ITS sequences of parents using PAUP* [32]. For that, we conducted a heuristic search with starting trees obtained via stepwise addition with 100 iterations of the random addition sequence and the TBR branch-swapping option in PAUP*. All constants and variable (i.e. non-informative) characters were deleted. Only informative characters for parsimony were examined. Consistency index (CI) and retention index (RI) were calculated and number of character changes mapped on branches. Then, for each putative hybrid, we built an MP unrooted tree in combining its ITS sequences with those of parents using the same method of reconstruction and only parsimony-informative characters. Finally, a MP phylogenetic tree was built for all *trn*K*-mat*K sequences of putative hybrids, *C. cordata* var*. cordata*, *C. griffithii*, meanwhile *C*. *nurii* var. *nurii* and *C*. *schulzei* as outgroups.

## Results

### Sequence analysis of nrITS region

The aligned length of nuclear ITS sequences had a total 736 bp with indicated twelve fixed nucleotide substitutions and indels which distinguished the *C. cordata* var*. cordata* sequences from the *C. griffithii* sequences at species level (Table 2). All fixed nucleotide substitutions with all clone accession numbers were summarized in S1 Table. For the putative hybrid, all individuals showed chromatogram peak additivity at all these fixed sites. No intraspecific polymorphism was detected within each putative parental species. Among the 42 cloned sequences of *C. ×purpurea* nothovar. *purpurea*, fourteen haplotypes (H1) were identical to *C. griffithii;* six haplotypes (H2) were identical to *C. cordata* var*. cordata*, and the remaining twenty-two cloned sequences (H3–H7) showed intermediate sequences between *C. cordata* var*. cordata* and *C. griffithii*. Of the six cloned sequences from the *C. cordata* var*. cordata* and *C. griffithii* template mixture, one was pure *C. griffithii* (H1), one was pure *C. cordata* var*. cordata* (H2), and the remaining four revealed intermediate sequences (H3). On the other hand, *C*. *nurii* var. *nurii* had ITS sequences different from those of *C. ×purpurea* nothovar. *purpurea* at six positions which eliminates *C*. *nurii* var. *nurii* as a possible parent. However, *C*. *schulzei* had identical ITS profiles to those of *C. cordata* var*. cordata* and *C*. × *purpurea* nothovar. purpurea. This additivity strongly supports *C. ×purpurea* nothovar. *purpurea* being the hybrid of *C. cordata* var*. cordata* and *C. griffithii*, although ITS data alone cannot reject the possibility of being *C. griffithii* × *C*. *schulzei*.

**Table 2.**
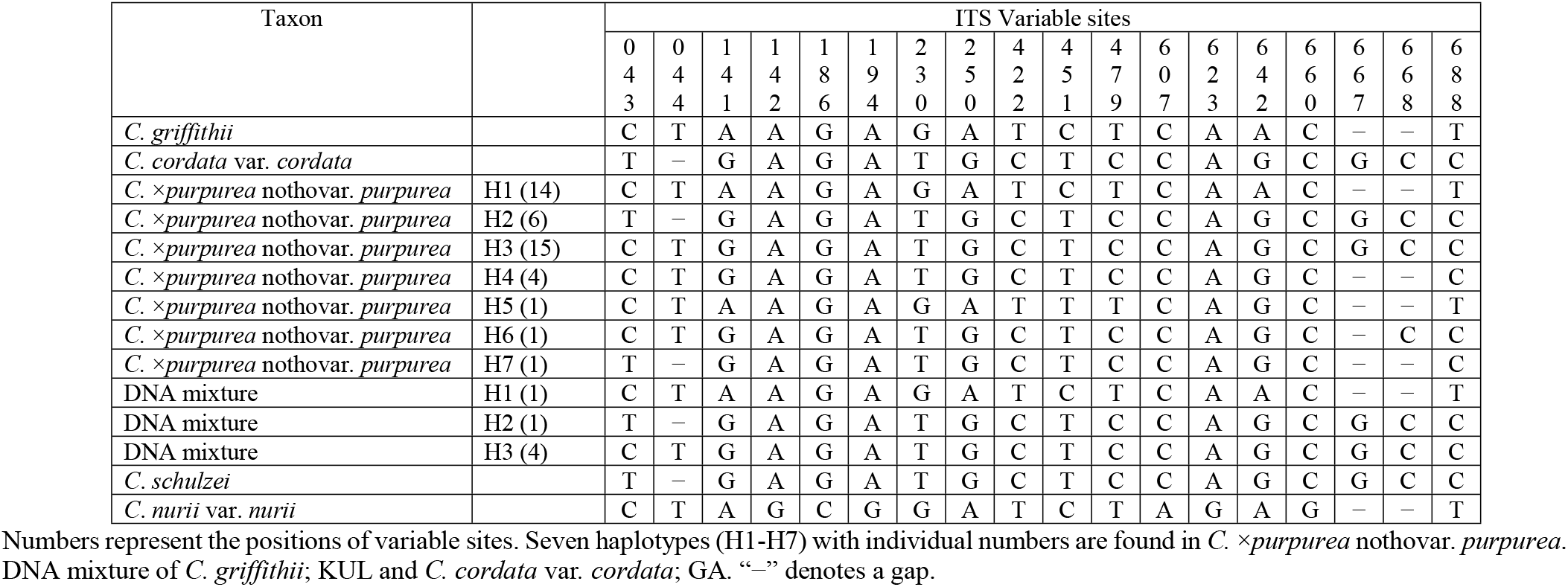
Variable nucleotide sites of ITS sequences comparison between the clones and the putative parental species.

MP analysis based on equal weighting of each character, yielded one tree of twelve steps, with a CI of 1 and RI of 1. Fig 2A showed this tree in which the two putative parents, *C. griffithii* and *C. cordata* var. *cordata*, form two distinct groups that differ to each other by nine unique substitutions (=fixed differences). Analysis of the putative clonal hybrid sequences showed that, for four putative hybrids (PK, MT, KJ and SL), we identified two groups of sequences corresponding to *C. cordata* var. *cordata* and *C. griffithii*. These two groups differ from each other by six to eight unique substitutions. For putative hybrids PI, SED and SU, only sequences similar to *C. cordata* var. *cordata* were detected from the six sequenced clones.

**Fig 2.**
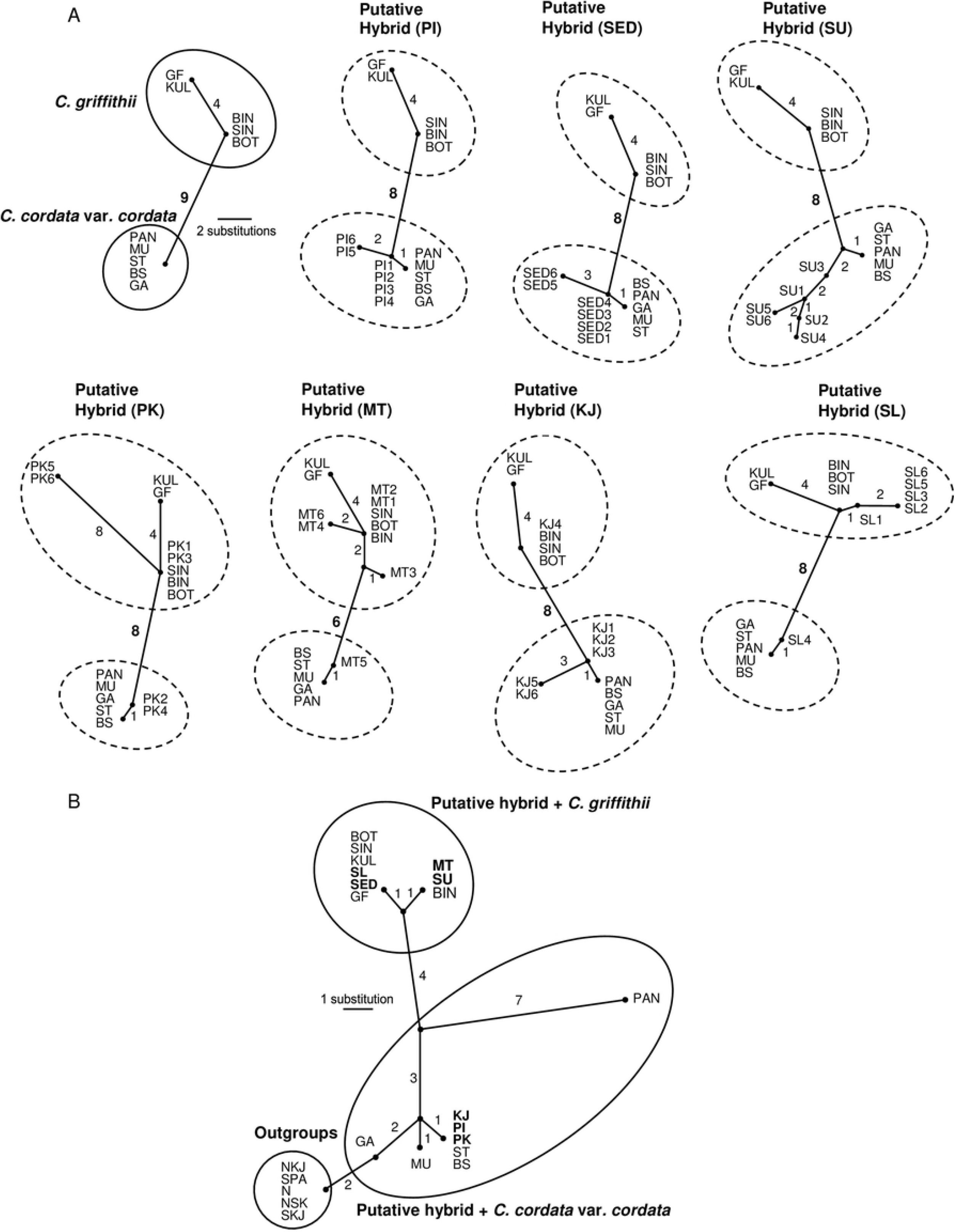
Most parsimonious (MP) unrooted tree based on equal weighting of each character based on ITS (A) and *trnK-matK* (B) sequences. Clone accession numbers were included in the ITS tree as per S1 Table. The numbers close to the connecting lines denote mutational steps.

### Sequence analysis of cpDNA region

The aligned length of chloroplast *trn*K-*mat*K region were 1980 bp long in the whole data set. An alignment of consensus nucleotide sequences from all samples varied at twenty sites (Table 3). For these positions, four substitutions and six single-base pair indels distinguished the *C. cordata* var*. cordata* sequences from the *C. griffithii* sequences. No variation was identified in species sampled from the same location. The comparison showed that the putative hybrid samples from PI, KJ and PK had sequences identical to *C. cordata* var*. cordata* of ST and BS meanwhile SL and SED had sequences identical to *C. griffithii* (GF, KUL, BOT, SIN). The *C. ×purpurea* nothovar. *purpurea* accessions from MT and SU had sequences identical to *C. griffithii* (BIN) at one nucleotide position. The *trnK-matK* sequences of *C. cordata* var*. cordata* (PAN) differed by three substitutions and *C. cordata* var*. cordata* (GA) differed by two substitutions when compared to other *C. cordata* var*. cordata* accessions. The nucleotide composition for *C*. *nurii* var. *nurii* is identical to *C*. *schulzei* and dissimilar to those of *C. ×purpurea* nothovar. *purpurea* at two nucleotide positions which makes it unlikely that both *C*. *nurii* var. *nurii* and *C*. *schulzei* can be the parents to *C*. × *purpurea* nothovar. *purpurea*.

**Table 3.** Variable nucleotide sites of *trnK-matK* sequences comparison between *C. ×purpurea* nothovar. *purpurea* and the putative maternal parents.

| Taxon | Variable sites |  |  |  |  |  |  |  |  |  |  |  |  |  |  |  |  |  |  |  |
| --- | --- | --- | --- | --- | --- | --- | --- | --- | --- | --- | --- | --- | --- | --- | --- | --- | --- | --- | --- | --- |
|  | 0 | 0 | 0 | 0 | 0 | 0 | 0 | 0 | 0 | 0 | 0 | 0 | 0 | 0 | 1 | 1 | 1 | 1 | 1 | 1 |
|  | 1 | 1 | 1 | 1 | 1 | 1 | 1 | 2 | 2 | 3 | 4 | 4 | 5 | 5 | 0 | 0 | 1 | 1 | 2 | 3 |
|  | 4 | 6 | 6 | 6 | 6 | 6 | 6 | 4 | 6 | 6 | 1 | 9 | 4 | 5 | 5 | 8 | 7 | 8 | 3 | 8 |
|  | 8 | 4 | 5 | 6 | 7 | 8 | 9 | 1 | 3 | 7 | 0 | 4 | 3 | 0 | 0 | 9 | 6 | 2 | 9 | 9 |
| *C. ×purpurea* nothovar. *purpurea*; PI | G | — | — | — | — | — | — | T | T | G | T | T | T | T | G | A | T | T | C | T |
| *C. ×purpurea* nothovar. *purpurea*; KJ | G | — | — | — | — | — | — | T | T | G | T | T | T | T | G | A | T | T | C | T |
| *C. ×purpurea* nothovar. *purpurea*; PK | G | — | — | — | — | — | — | T | T | G | T | T | T | T | G | A | T | T | C | T |
| *C. cordata* var. *cordata*; ST | G | — | — | — | — | — | — | T | T | G | T | T | T | T | G | A | T | T | C | T |
| *C. cordata* var. *cordata*; BS | G | — | — | — | — | — | — | T | T | G | T | T | T | T | G | A | T | T | C | T |
| *C. cordata* var. *cordata*; MU | G | — | — | — | — | — | — | T | T | G | T | T | G | T | G | A | T | T | C | T |
| *C. cordata* var. *cordata*; GA | G | — | — | — | — | — | — | T | T | T | T | T | G | T | G | A | T | T | C | A |
| *C. cordata* var. *cordata*; PAN | G | — | — | — | — | — | — | T | T | G | C | T | G | T | A | A | G | T | C | T |
| *C. ×purpurea* nothovar. *purpurea*; SL | G | C | T | G | T | A | T | T | G | G | C | T | G | G | A | G | G | C | A | T |
| *C. ×purpurea* nothovar. *purpurea*; SED | G | C | T | G | T | A | T | T | G | G | C | T | G | G | A | G | G | C | A | T |
| *C. griffithii*; GF | G | C | T | G | T | A | T | T | G | G | C | T | G | G | A | G | G | C | A | T |
| *C. griffithii*; KUL | G | C | T | G | T | A | T | T | G | G | C | T | G | G | A | G | G | C | A | T |
| *C. griffithii*; BOT | G | C | T | G | T | A | T | T | G | G | C | T | G | G | A | G | G | C | A | T |
| *C. griffithii*; SIN | G | C | T | G | T | A | T | T | G | G | C | T | G | G | A | G | G | C | A | T |
| *C. ×purpurea* nothovar. *purpurea*; MT | A | C | T | G | T | A | T | T | G | G | C | T | G | T | A | G | G | C | A | T |
| *C. ×purpurea* nothovar. *purpurea*; SU | A | C | T | G | T | A | T | T | G | G | C | T | G | T | A | G | G | C | A | T |
| *C. griffithii*; BIN | A | C | T | G | T | A | T | T | G | G | C | T | G | T | A | G | G | C | A | T |
| *C. schulzei*; SPAN | G | — | — | — | — | — | — | G | T | T | T | A | G | T | G | A | T | T | C | A |
| *C. schulzei*; SKJ | G | — | — | — | — | — | — | G | T | T | T | A | G | T | G | A | T | T | C | A |
| *C. nurii* var. *nurii*; NKJ | G | — | — | — | — | — | — | G | T | T | T | A | G | T | G | A | T | T | C | A |
| *C. nurii* var. *nurii*; NSK | G | — | — | — | — | — | — | G | T | T | T | A | G | T | G | A | T | T | C | A |

Numbers represent the positions of variable sites. “—” denotes a gap.

Phylogenetically informative under parsimony based on equal weighting of each character and including outgroup sequences, yields one tree of 20 steps, with a CI of 1 and RI of 1. In this tree (Fig 2B), the ingroup and *C. griffithii* are each monophyletic, while *C. cordata* var. *cordata* is not monophyletic because outgroups are nested within. When the *trn*K*-mat*K sequences of the putative hybrids are added to the dataset, the resulting MP tree shows that the maternal origin of the *trnK-matK* sequences of the putative hybrids KJ, PI and PK is *C. cordata* var. *cordata* whereas *C. griffithii* is the parent of the *trn*K*-mat*K sequences of the putative hybrids SL, SED, MT and SU.

## Discussion

Natural hybridization has been considered to represent an important factor influencing the high diversity of the genus *Cryptocoryne* [7]. More *Cryptocoryne* species may frequently be observed in co-existence, inhabiting the same or adjacent streams which may lead to hybridization and the production of hybrid populations within the same area [6–10]. The capability of *Cryptocoryne* to hybridize naturally may create more complexity in terms of taxonomic studies and classification. Previous studies of hybridization in *Cryptocoryne* were based on morphology, cytology and crossing experiments [6–8, 20, 21], while the use of molecular markers in hybrid investigation within *Cryptocoryne* is almost absent.

The nuclear genes are biparentally inherited, and the hybrids should possess both divergent copies of their putative parents [26, 27]. The nuclear ITS sequences of *C. ×purpurea* nothovar. *purpurea* showed nucleotide polymorphism at some sites whereas those of *C. cordata* var*. cordata* and *C. griffithii* are not; the data from *C*. × *purpurea* nothovar. *purpurea* taxa exhibited polymorphism patterns which is consistently formed by additive sequences derived from the two hypothesized parental species. The present ITS data revealed that *C. ×purpurea* nothovar. *purpurea* possessed hybrid genotypes; having both ITS haplotypes of parental *C. cordata* var*. cordata* and *C. griffithii* suggests that *C*. × *purpurea* nothovar. *purpurea* is a natural hybrid between these two species. The results also showed that 22 (52.4%) out of the 42 cloned nrITS sequences from *C. ×purpurea* nothovar. *purpurea* were intermediate/chimeric (recombinations of parental sequence haplotypes (H3-H7)). In diploid hybrids, the co-occurrence of parental nrITS sequences is rarely maintained in subsequent generations due to concerted evolution; if concerted evolution is incomplete, then sampled genes may represent a mixture of non-homogenized paralogous sequences [33]. Effects of concerted evolution commonly occurs after meiosis (sexual reproduction) only in fertile plants [34, 35]. However, Jacobsen [14] reported that the pollen of *C. ×purpurea* nothovar.*purpurea* is completely sterile. Cytological analysis shows that *C*. ×*purpurea* nothovar. *purpurea* shares the same diploid chromosome number 2*n* = 34 as *C. cordata* var*. cordata* and *C. griffithii*, which rules out the possibility of *C*. × *purpurea* nothovar. *purpurea* being a sterile polyploid. Because *C*. × *purpurea* nothovar. *purpurea* is sterile, all *C*. × *purpurea* nothovar. *purpurea* individuals are assumed to represent an F_1_ generation. Therefore, F_1_ individuals should possess both parental ITS alleles without impact of concerted evolution. One explanation for the origin of such unique allele in *C*. × *purpurea* nothovar. *purpurea* may be the result of PCR-mediated recombination, a process of *in vitro* chimera formation from related DNA template sequences coamplified in a single PCR reaction [36]. PCR-mediated recombination results from either polymerase template switching during PCR or annealing of prematurely terminated products to non-homologous templates [36, 37] suggesting that the chimeric ITS sequences found in *C*. × *purpurea* nothovar. *purpurea* can be due to artificial recombinants. Moreover, the chimeric haplotypes are unequally distributed in the nucleotide positions which indicates the process of recombination occurring in a non-random order during PCR. Techniques involving passing traditional PCR and cloning process are necessary for deeper examination of the structure and evolution of nrITS sequences in *C*. ×*purpurea* nothovar. *purpurea*. Due to the high sterility of *C*. × *purpurea* nothovar. *purpurea*, derived from the low pollen stainability with cotton blue [12], it may be inferred that sterility is also prevalent of the female side. However, we cannot exclude that a backcrossing from one of the putative parents may take place, but we have not observed any cases where that might be the case in the present context, or maybe our material is too limited to draw such conclusions. In other cases, regarding *Cryptocoryne* species from Sri Lanka [18] second or more generations have been reported and assumed, but here the hybrids showed some degree of fertility. In the situation regarding *C*. *crispatula* Engl. *s.l*. multiple hybrids have been observed and there is some fertility in the hybrids, and backcrosses and/or introgressions is suggested to be highly likely [7].

The results showed that the ITS sequences of *C. cordata* var*. cordata* and *C*. *schulzei* are very similar suggesting that the *C. ×purpurea* nothovar. *purpurea* populations examined are hybrids between *C. griffithii* and either *C. cordata* var*. cordata* or *C*. *schulzei*. The hybrid grows sympatrically with *C. griffithii* and *C. cordata* var*. cordata* and shows obvious morphological intermediary between the two species. *Cryptocoryne* species are mainly identified using floral characters, particularly the limb of the spathe. The limb of the spathe of *C*. × *purpurea* nothovar. *purpurea* provides strong evidence in identification of the parental species by showing intermediate characters to the two parents; - the broad collar zone being present in both *C*. × *purpurea* nothovar. *purpurea* and *C. cordata* var*. cordata*; but a rather rugose limb of the spathe with wide pronounced collar is clearly visible in *C*. *schulzei*. A rough purple red limb of the spathe is characteristic of *C. griffithii* and resembles that of *C. ×purpurea* nothovar. *purpurea*; *C*. *nurii* var. *nurii* has a deep red to dark purple spathe with large irregular protuberances on the limb. In conclusion, with the joint examination of molecular and morphological datasets from included accessions, it is unlikely that *C*. *schulzei* and *C*. *nurii* var. *nurii* are parents to *C*. ×*purpurea* nothovar. *purpurea*. In the above-mentioned study on artificial hybridization between species of *Cryptocoryne* from the Malay Peninsula, hybrids were produced between *C. cordata* var. *cordata* and *C. nurii* var. *nurii* which were not like *C. ×purpurea* nothovar. *purpurea* [8].

It is common to find more than one species of *Cryptocoryne* in southern Peninsular Malaysia (Pahang, Johor and Malacca) that share the same stream or river system. Suitable combinations of species coexist, resulting in hybridization conditions. The *trn*K*-mat*K sequences of *C*. ×*purpurea* nothovar. *purpurea* accessions PI, KJ and PK from Pahang (all originating from the Tasik Bera (Bera Lake) region) were identical to *C. cordata* var*. cordata* and pointed out this species as the maternal parent. Neither putative parent was found in the hybrid sampling lake, but *C. cordata* var*. cordata* (ST) was present in the nearby swamp. The origin of the Tasik Bera area dates back to only 4500 years B.P [38]. Based on the explanation of Othman et al. [6], the main drainage of the Tasik Bera is now northwards to Sungai Pahang (Pahang River), but there is still a small connection southward to the Sungai Palong/Sungai Muar, Johor that was formally the main run-off. This historical event provides the explanation of *C*. × *purpurea* nothovar. *purpurea* having arisen as a hybrid between more widespread *C. cordata* var*. cordata* and the southerly distributed *C. griffithii*, which has then spread along the west coast during the change in drainage systems. The *trnK-matK* sequences of *C. ×purpurea* nothovar. *purpurea* accessions from Johor (SL and SED) were identical to the *C. griffithii* populations from Johor (KUL and GF) and *C. griffithii* from Singapore (BOT and SIN). Currently it is known that the two putative parental species have both recently been found at the Sg. Sedili Kechil, Johor and previous records showed that *C. cordata* var*. cordata* and *C. griffithii* distribution overlapped in Johor [7]. The *C. griffithii* was also proposed as the maternal parent to the *C*. × *purpurea* nothovar. *purpurea* populations from Malacca region (MT and SU), viz. *C. griffithii* from BIN (Bintan, Indonesia). The other parental species *C. cordata* var*. cordata* (BS) were found within a distance of <40 km from all the hybrid locations in the Malacca region but *C. griffithii* has not been recorded from there recently. However, *C. griffithii* has previously been recorded to grow in several places in Malacca [6, 19]. The present results indicate that both *C. cordata* var*. cordata* and *C. griffithii* have served as maternal donor and the different hybrid populations possess separate and independent origins. There was no distinct bias of maternal composition for either one of them, and this suggests that natural hybridization between the two examined species is bidirectional.

The combined investigation of nuclear ribosomal DNA, viz. the ITS and the chloroplast DNA for the *trn*K-*mat*K region provide compelling evidence for the natural hybridization between *C. cordata* var*. cordata* and *C. griffithii* and the hybrid origin of the tested individuals. Molecular data supports the hypothesis that the morphologically intermediate plants are hybrids which share *trn*K*-mat*K sequences identical to both *C. cordata* var. *cordata* and *C. griffithii*. As *C*. × *purpurea* nothovar. *purpurea* is pollen sterile (and is also believed to be ovule sterile) with the absence of meiotic recombination, the parental sequences of the nuclear and chloroplast markers may be retained in the vegetative progenies. This study provides substantial evidence for interspecific hybridizations in *Cryptocoryne*. It should be interesting to further investigate the population genetics, ploidy level and reproductive behaviour of the hybrids including the geographical distribution and the timing for hybridization events for a better understanding of the total extent of hybridization process in *Cryptocoryne*.

## Acknowledgments

Sebastien Lavoue and Siti Nurfazilah are thanked for their useful comments on an earlier draft of this manuscript.

## Supporting information

**S1 Fig. The geographical distributions of *C. ×purpurea* nothovar. *purpurea* and the putative parental species**.

**S1 Table. Variable sites of ITS between the clone accession numbers and the putative parental species.**

